# Efficient and reproducible somatic embryogenesis and micro propagation in tomato via novel structures -Rhizoid Tubers

**DOI:** 10.1101/446286

**Authors:** Wajeeha Saeed, Saadia Naseem, Daniyal Gohar, Zahid Ali

## Abstract

An improved and highly reproducible system for *invitro* regeneration via somatic embryogenesis (S.E), applicable to several varieties of tomato (cv. *Riogrande*, cv. *Roma grande*, hybrid *17905* and model cv. M82) has been developed. First, we developed a conventional indirect organogenesis for all four varieties used in this study. The cotyledons and hypocotyls of 6-day-old tomato were used as explants (1-2 cm) for callus induction (CI) on different callus induction media (CIM) T_0_ – T_12_ (6-Benzylaminopurine BAP, NAA Naphthalene acetic acid, ZEA Zeatin, IAA Indole-3-acetic acid, KIN Kinetin). Maximum CI response was seen on CIMT_6_ (0.5 mg/L NAA, 1 mg/L BAP) and CIMT_7_ (2 mg/L IAA, 2 mg/L NAA, 2 mg/L BAP, 4mg/L KIN) in a period of 2 weeks for commercial varieties *cvs. Riogrande* and *Roma*. However, *cv. M82* responded after 4 weeks to a combination of treatments [CIMT_6_ (0.5 mg/L NAA + 1 mg/L BAP) and CIMT_8_ (2 mg/L IAA + 2 mg/L NAA + 2 mg/L BAP + 4 mg/L ZEA)] for the production of calli. The *Riogrande*, being the most responsive commercial variety, was selected for *invitro* morphogenesis via S.E. During S.E. young cotyledons and hypocotyls explants were tested on media with different ranges of pH (3 – 7) supplemented with 0.5 and 2 mg/L NAA. Resultantly, numerous rhizoids (~38) were produced from each explant at pH4 in dark conditions. Further incubation of each rhizoid under light conditions led to the formation of a novel structure - rhizoid tubers (RTBs) on MS media supplemented with 5 mg/L TDZ/BAP at pH4. We observed that only lower pH-induced rhizoids and RTBs regenerated into multiple individual shoots on media at normal pH (5.8). The RTBs led to a complete plantlets regeneration in 45 days compared to the conventional *invitro* morphogenesis (60 days).

## Introduction

Tomato (*Solanum lycopersicum*) is the most important edible perennial crop from the *Solanaceae* family. In addition to its high nutritious value, tomato also serves as a great tool for advanced biotech research. Its relatively smaller genome, a diverse germplasm, and the availability of a suitable transformation system for it make tomato an ideal and convenient model system for the improvement of other dicotyledonous plants [1,2]. About 177 million tons of tomatoes were produced in 2016 worldwide [3]. A quarter of this produce is being processed in industry, thus making this commodity the world’s leading vegetable processing industry. China is playing a leading role in tomato production followed by India and United states. In the Euro zone, about 18 Million tons of tomatoes were produced in 2016 of which 2/3 came from Italy and Spain [4]. In Pakistan, about 0.6 Million tons of tomatoes have been produced from 62,930 hectares of land in 2014 [5]. Despite a favorable environment and an available infrastructure for the mass production of tomato, marketable yields have been declining due to environmental constraints and pre- and post-harvest losses [6]. Being a highly perishable horticulture, tomatoes are prone to post-harvest losses. These losses, however, are more prominent in the developing countries where loss index reaches up to 50% annually [7,8]. Several other biotic and abiotic factors also reduce the annual production of tomato globally. Sustainable tomato cultivation requires conventional breeding practices and crop management to overcome these barriers.

There is an increased demand for the introduction of qualitative traits into commercially important cultivars of tomato in order to increase productivity, ameliorate nutritional value, and extend shelf life. [1,9,10]. Conventional breeding and micropropagation and/or genetic engineering have become valuable tools for the incorporation of important traits. *Invitro* micro propagation and plant cell cultures offer a tremendous prospect of utilizing cell/tissue and organ culture for genetic improvement of important horticultural crops. A high throughput as well as a reproducible *invitro* regeneration system that leads to time-efficient recovery of transgenics is inevitable. Most often, the morphogenesis and totipotency of tomato have been reported to be lower than that of the other members of *Solanaceae* [11,12]. In tomato, the effect of genotype is most significant, and one set of plant hormones that leads to regeneration might not work for other closely related cultivars, hence imposing laborious constrains and prolonged regeneration responses. Different explants used for micropropagation and/or transformation in various model cultivars from leaf [13], cotyledons [14,15], hypocotyls [16], meristems [17], inflorescence [18], anthers [19], suspension cells [20] have also highlighted intractability of regeneration in tomato through organogenesis. Nevertheless, regeneration in tomato could be achieved via organogenesis and direct or indirect somatic embryogenesis (S.E) from organized tissues. Direct S.E has a favorable outcome for an invitro regeneration response and a highly efficient transformation system with cytological fidelity in a short time whereas reports to achieve regeneration via S.E are limited [21].

Although S.E has the potential to produce a large no. of elite/transgenic plantlets, its possible application in tomato is still in infancy. In this regard, considerable advancements can uplift the use of S.E for *in vitro* propagation as well as genetic improvement in tomato. The current study is aimed at the development of a simple yet robust *invitro* regeneration of four different cultivars [Two commercial cultivars *cv*. *Riogrande, Romagrande*, one local hybrid-17905 and *M82* (model cultivar)] through organogenesis and S.E. Different explants sources were compared for *invitro* regeneration frequency. The effect of genotype and growth regulators on a systemic survival of regenerated plants was accessed. We present a new and improved regeneration system via S.E involving rhizoid tubes (RTBs) which were developed through a novel approach. Furthermore, it is proposed that regeneration via S.E and RTBs can be utilized for the development of transgenic plants with high cytological fidelity.

## Materials and methods

### Plant material and seed disinfection

The research was carried out at Plant Biotechnology & Molecular Pharming (PBMP) Lab, COMSATS University Islamabad, Pakistan during 2016-2018. The seeds of three local tomato cultivars *cv. Riogrande, cv Romagrande*, Hybrid-17905 (from Plant Genetic Resources. NARC-Islamabad), and one model *cv. M82* were used. Seed storage was maintained at 4°C in the dark. For seed sterilization, the seeds were first rinsed with autoclaved, double-distilled water followed by immersion in 70% ethanol for 1-2 minutes and then transferred to four different combinations of sterilizations [(NaOCl), (v/v1%-20%), 6% NaOCl with 2 drops of tween 20 and house hold bleach (8%)] (Supplementary material Table S1). Following sterilization, the seeds were washed five times with deionized autoclaved water and blotted dry on sterilized filter paper. 150 seeds for each treatment were utilized in three replicates. The seeds were aseptically placed on MS growth media plus vitamins [22] full strength and half strength, with and without 2% sucrose solidified with 4% phytagel and 8% plant agar. All basal media consisted of MS with vitamins 4.15 g/L with 8% agar and 20 g sucrose (**Table 1**). The pH of the medium was adjusted to 5.8 before autoclaving. The seeds were kept for 48hrs in the dark and subsequently maintained at 23°C±2, with 30–50% humidity and 16/8hrs Light/Dark photoperiod provided by 70 μmolm^-2^s^-1^ cool white fluorescent lights in a growth room. Germination index for the newly emerged seedling was calculated from the age of 8 days until 80% germination was achieved.

**Table 1.** Optimized media formulations used in Tomato invitro morphogenesis and transformation

| Optimized medium used for regeneration and transformation of tomato |  |
| --- | --- |
| Culture medium | Additional Components |
| Germination medium (GM) | 3.17g/L MS salts, 8g/L plant Agar and pH 5.8 |
| Callus induction medium (CIM) | 2mg/L NAA, 2mg/L IAA, 2mg/L BAP, 4mg/L KIN or 4mg/L Zeatin, phytigel 4g/L and pH 5.8 (Cv. M82) |
| Shoot induction Medium (SIM) | MS salts, 20g/L sucrose, 3mg/L BAP and 0.1mg/L IAA |
| Root induction Medium (RIM) | MS salts, 20g/L sucrose and 0.5mg/L NAA or 1mg/L IBA |
| Rhizoids induction medium (RhIM) | MS salts, 10g/L sucrose, 0.5 or 2 mg/L NAA and pH4 |
| Tubers induction Medium (TIM) | MS salts, 10g/L sucrose, 5mg/L BAP or 5mg/L TDZ and pH4 |
| Pre-Culture Medium (PCM) | MS salts, 20g/L sucrose, 1mg/L NAA and BAP 1mg/L |
| Co-Cultivation Medium (CCM) | MS salts, 20g/L sucrose and 200 μM *Acetosyringone*, |
| Selection Medium (SM) | MS salts, 20g/L sucrose, 2mg/L IAA, 2mg/L BAP, 4mg/L KIN or 4mg/L, 300mg/L cefotaxime/ 300mg/L Augmentin, and 600 mg/L ticarcillin |
| Infiltration Medium (IFM) | MS salts, 20g/L sucrose, 2mg/L NAA and 100 μM *Acetosyringone* |
| Inoculation Medium (IM) | MS salts, 20g/ L sucrose and 200 μM *Acetosyringone* |

### Callus induction on different media formulations

Following the germination, 6- to 7-days old young cotyledons, hypocotyls, and a stem section were cut aseptically. Curled cotyledons were cut from both ends to form a 0.5−1 cm long leaf disc. Leaf discs were placed abaxial side down. The stem section and hypocotyls were also placed horizontally on a callus induction medium (CIM). MS salts with vitamins [22] was used as the basal medium with 10,15,30 g/L sucrose, 8% plant agar and/or 4% phytagel and pH 5.8. Different plant growth hormones (PGRs) Naphthalene acetic acid (NAA), 6-Benzylaminopurine (BAP), Indole-3-acetic acid (IAA), Kinetin (KIN), Zeatin (ZEA), 2,4-Dichlorophenoxy acetic acid (2,4-D) and Gibberellic acid (GA3) were used alone and in combination viz: CIMT_0_-CIMT_12_ (Table 2). Percent callus induction on different hormonal combinations repeated in triplicate was recorded after 4 weeks of induction. Callus morphology and regeneration were also observed.

**Table 2.** The Effect of different concentrations and combinations of PGRs with MS medium on callogenesis and regeneration of four tomato varieties

| Treatment (Callus induction Medium) CIM | ROMA | Rio Grande | 17905 | M82 |
| --- | --- | --- | --- | --- |
| CIMT<sub>0</sub>=MS salts+vitamins 3%Suc+8g/L agar | – | S- |  | – |
| CIMT<sub>0</sub>1=1mg/L NAA+1mg/L BAP | CR (++++) | CR (++) | CR (++) | C (+) |
| CIMT1= 0.2-2 mg/L NAA | R<sub>1</sub> (+++) | R<sub>1</sub> (+++) | R<sub>1</sub> (+++) | R<sub>1</sub> (++) |
| CIMT2= 0.2mg/L NAA+1mg/L BAP | CR (++) | CR (+) | CR (++) | C (+) |
| CIMT3=0.2mg/L NAA+2mg/L BAP | CR (+) | RS (+) | RS (+) | S (+) |
| CIMT4=0.2mg/L NAA+3mg/L BAP | CSR (+) | RS (+) | S (+) | S (+) |
| CIMT5=0.2mg/LNAA+4mg/LBAP | CS (+) | S (+) | S (+) | S (+) |
| CIMT6= 0.5 mg/L NAA+ 1mg/L BAP | C (++++) | C (+++++) | C (+++) | C (+++) |
| CIMT7=2mg/L IAA+2mg/L NAA+2mg/L BAP +4mg/L Kin | C (+++++) | C (+++++) | C (++++) | C (+) |
| CIMT8=2mg/L IAA+2mg/L NAA+2mg/L BAP +4mg/L Zea | - | - | - | C (++++) |
| CIMT9=0.5mg/L IAA+2-0.5mg/L NAA+2mg/L 2,4-D+0.2mg/Lm Zea | - | - | - | CR (++++) |
| CIMT10=1=0.5mg/L IAA+1mg/L BAP | C (++) | C (++) | C (++) | C (+) |
| CIMT11=0.5mg/L BAP+0.5mg/L NAA+2mg/L GA3 | C (+++) | C (+++) | C (+++) | - |
| CIMT12=2mg/L 2,4-D+0.5mg/L BAP | - | - | - | C (+) |

| Rating Scale |  | Comments |
| --- | --- | --- |
| Rating | Callusing index |  |
| +++++ | >70% & ≤ 80% | R: Regeneration |
| ++++ | ≤ 70% | R<sub>1</sub>: Rooting |
| +++ | ≤50%but >30% | S: Shooting |
| ++ | >20% | C: Callus formation |
| + | <10 | CSR: Callus leading to regeneration |
| - | 0% | CR: Callus with rooting |

### The Effect of pH on callus induction and rhizoids tubers

For the optimization of somatic embryogenesis from leaf and cotyledon explants, supplementary auxin concentration and medium pH were tested. The effect of pH on the overall growth of the callus and rhizoid induction was evaluated by the media formulation (MS salts with vitamins 4.13 g/L, sucrose 3% and agar 0.8%) with the pH ranges of (3, 4, 5, 5.8, 6, 7) supplemented with NAA (at 0.5, 1, 1.5, 2 and 4 mg/L) and 2,4-D (at 2, 3 and 5 mg/L). The optimization of rhizoids and the formation of rhizoids tubers (RTBs) from cotyledons and hypocotyl explants were done in the dark at 23°C ± 2 with an optimal concentration of two auxins analogues and a series of medium pH. After a considerable number of rhizoids formed, further sub-culturing of each rhizoid was carried out under 16/8hrs photoperiod (Light/Dark) with 5 mg/L TDZ and/or BAP on pH ranges (3–7) along with pH 5.8 to test RTBs formation. The rhizoids formed per explant and their invitro morphogenesis were counted and stages of RTBs formation were recorded by using Nikon D5200 Camera.

### Microscopic Investigations

For histochemical staining, RTBs were fixed and dehydrated by using Formalin:Glacial acetic acid:Ethanol as a fixing solution [23]. Paraffin-embedded samples were cut into 12 mm thick transverse sections using rotary Microtome (Amos scientific AEM 480) and stained with 1% safranin stain. The sections were observed under an Automatic scanning system (Zeiss AS3000B - UK) and stereomicroscope and respective images were captured at different magnifications (25–100X) as per requirement.

### Plantlet regeneration from callus and RTBs

A healthy, proliferating green/pale yellow callus was sub-cultured routinely. The calli formation and RTBs developed from cotyledons were evaluated on different shoot-induction media (SIM) viz SIM_1_–SIM_6_ (Table 5) to compare the organogenesis response. The effect of BAP and TDZ (2–5 mg/L) with and without auxins was investigated for a maximum number of regenerated shoots from callus and RTBs. For a standard shoot-induction response MS salts (w/vitamins 4.13 g/L, 2% sucrose, 0.8% agar and pH 5.8) was used. Alongside, pH range (4–7) was tested for in vitro RTBs on MS media supplemented with 3–5 mg/L BAP w/wo auxins. Individual calluses/calli and RTBs were placed on shoot induction media (SIM) with 16/8hrs Light/Dark photoperiod at 23°C± 2, under white cool fluorescent lights (70 μmolm^-2^s^-1^) and subcultured after every 3 weeks. The number of induced shoots was recorded 2 weeks after induction.

### Rooting and shoot elongation

3–6 weeks old *invitro* regenerated shoots (3–4 cm) from callus and individual rhizoid tubers were cut aseptically and transferred to a rooting medium containing IBA (0.1–1 mg/L), NAA (0.1–0.5 mg/L) and IAA (0.1/0.2 mg/L). The effect of PGRs on rhizogenesis was recorded on the rooting medium. Data was recorded as % number of individual shoots rooted on RIM (root induction medium).

### Ex-vitro acclimatization and transfer of rooted plants

Regenerated explants were grown on RIM for four weeks until they were 3–5 cm in length. Plants with fully grown roots were removed from the medium and gently washed with tap water to remove the extra medium. The plants were then transferred to sterile soil and vermiculite mixture (1:1) in plastic pots and covered with a clear plastic bag with holes to sustain humidity level. Plants hardened for one and a half month were then transferred to a greenhouse with normal daylight conditions for flowering and fruiting.

### Statistical analysis

The data was analyzed using the analysis of variance (ANOVA) via SPSS 23.0, with 99% and 95% confidence intervals.

## Results

In the present study, various physical and chemical factors were optimized for commercially important as well as model cultivars of tomato for somatic embryogenesis and invitro morphogenic regeneration.

### Seed germination and contamination control

The containment of various types of fungal or bacterial contamination during *invitro* micropropagation is one of the prime prerequisites of successful tissue culture. Seed sterilization and storage conditions affect the overall process of *invitro* morphogenesis. We evaluated the effect of commercial sodium hypochlorite with and without tween 20 and household bleach on seed germination and their efficiency to control contamination (see supplementary Table S1). We achieved contamination-free seed germination without sucrose in our experiments (Table 1). Although sucrose is an important component required for tomato cell cultures, an absence of sucrose also yielded seedling emergence in the present study.

### Invitro callus culture induction is genotype-dependent

Various Plant Growth Regulators (PGRs) regulate an *in vitro* morphogenesis response by modulating different physiological processes. In the current study, different combinations of PGRs have been tested for callus induction (CI) and regeneration. Two types of explants (cotyledons and hypocotyls) from 6–8 days old seedlings were used for callus induction which showed variable responses to different callus induction mediums/media (CIM) (**Table 2**). CI and regeneration from cotyledons and hypocotyls varied based on hormonal treatment and genotype. Maximum calli were developed from young cotyledons of *Riogrande*, having efficiency of 82.09% from CIMT_6_, 86% from CIMT_7_ and 72% from CIMT_11_. CIMT_7_ was found to be the most suitable treatment that leads to soft, fleshy green callus that quickly leads to regeneration. Roma showed callus formation on CIMT_7_ and CIMT_6_ being 85.9% and 75.89% respectively (**Table 2**).

However, the other two genotypes i.e. hybrid (*17905*) and *M82* were comparatively least responsive at above-mentioned hormonal treatments and thus took a longer time. In comparison model cultivar *M82* only exhibited callus induction activity against a combination of treatments (CIM T_6_ & CIM T_8_). Cotyledon discs were first placed on CIMT_6_ for 15 days after which swollen explants were sub-cultured on CIMT_8_. The CIMT_6_ or CIM_8_ alone were found ineffective in producing optimal callus formation. Rio and Roma showed healthy green friable callus proliferation on CIMT_7_ which were frequently sub-cultured for organogenesis. The 12 different treatments *viz* CIMT_0_-CIMT_12_ for efficient callus induction were highest to lowest in the order CIMT_7_>CIMT_6_>CIMT_01_>CIMT_11_. The highest callus induction was observed from the treatment CIMT_7_ having 2mg/L BAP, 2mg/L NAA, 2mg/L IAA and 4mg/L kinetin in both cotyledons and hypocotyls. Embryogenic calli derived from leafy explants depicted a high shoot regeneration potential in Rio and Roma while other two cultivars had pale white compact callus (Fig 1). A hypocotyl-derived callus had many embryoids in *Riogrande*, however, callus induction was slower, and calli were less totipotent as compared to cotyledons. The response to callogenesis was found highly genotype-dependent and was affected remarkably by the reproductive background (self-pollination) of the cultivars.

**Fig 1.**
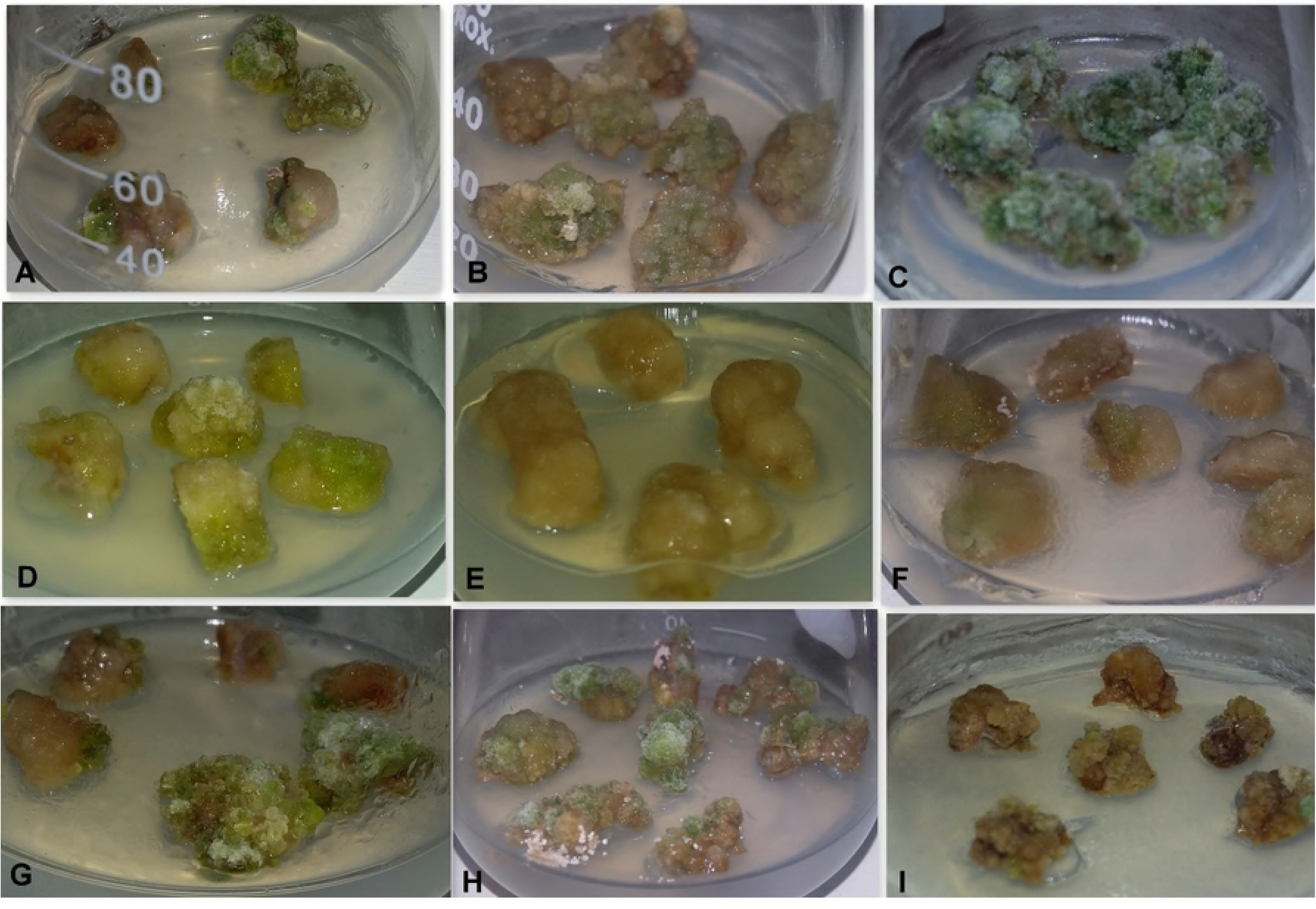
The Effect of different concentrations and combinations of PGRs with MS medium on callogenesis and regeneration of four Tomato varieties. (A, B) Calli induced from hypocotyls of Rio (C) Calli induced from leaf discs of Rio (D, E) Calli induced from leaf and hypocotyl of Roma (F, G) Calli induced from hypocotyls and leaf discs of Hybrid 17905 (H, I) Calli induced from leaf and hypocotyl of M82.

### Lower pH induced regeneration and RTBs

Lower pH medium supplemented with auxins induced somatic embryogenesis in 6–day-old tomato cotyledon and hypocotyl explants. The variable pH range (3–7) tested with two auxins (NAA and 2,4-D) with concentrations 0.5 mg/L and/or 2 mg/L revealed that only pH 4 favored rhizoid formation in dark conditions (Figs 2 and 3). Without PGRs and dark conditions no rhizoids were observed while pH 3 led to a failure in the solidification of the medium to support tubers or morphogenesis. Different concentrations of both auxins analogues tested in triplicate with all pH levels revealed that 2,4-D was not as effective as NAA [0.5 and 2 mg/L] (**Table 3**). The results showed that explants subcultured on a medium supplemented with 2,4-D failed to produce rhizoids.

**Fig 2.**
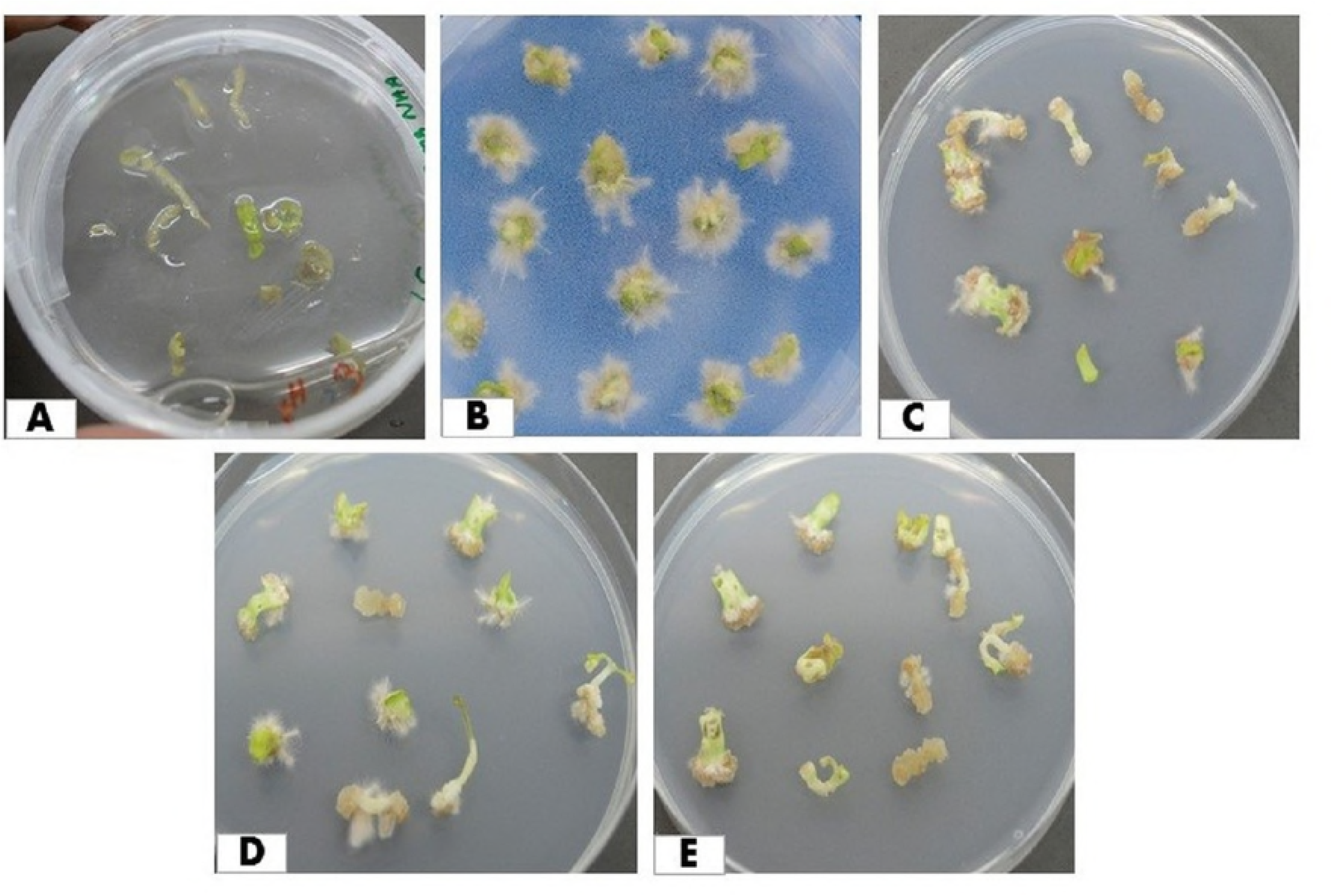
The effect of medium pH (3-7) on rhizoids production from cotyledon eplants of tomato (*cv Riogrande*) with optimized concentration of NAA (0.5/2 mg/L). (A) pH3, (B) pH4, (C) pH5, (D) pH6, (E) pH7.

**Fig 3.**
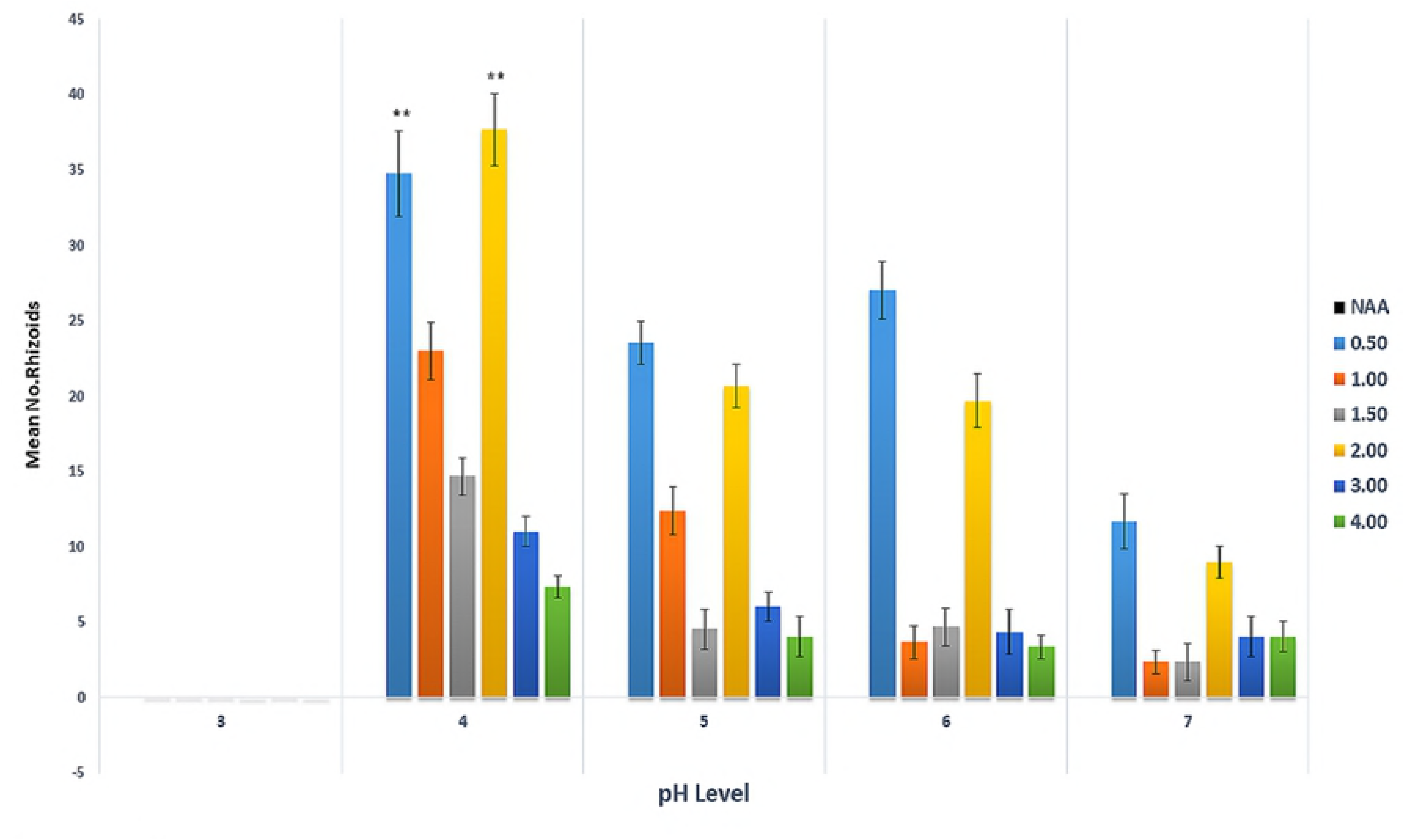
The effect of NAA concentrations against medium pH (3–7) for induction of rhizoids from cotyledon explants of tomato (*cv Riogrande*) Values are represented as mean ± standard error per treatment calculated from 90 explants each treatment. Significant differences were analyzed with Tukey’s-HSD test at *p* <0.001

**Table 3.** The Effect of Various Concentrations of NAA on Rhizoids Formation from Cotyledon/Cypocotyl Explants at pH4.

| NAA (mg/L) | Average number of Rhizoids per Ex-Plant | 2,4-D (mg/L) | Average numbers of Rhizoids per explant at |
| --- | --- | --- | --- |
| 0 | 0.00±0.00 | 0 |  |
| 0.5 | 38.33± 2.60** | 0.5 |  |
| 1 | 23± 2.08 | 1 |  |
| 1.5 | 14.66± 0.88 | 1.5 | 0.00±0.00 |
| 2 | 37.667± 3.28** | 2 |  |
| 3 | 11± 0.57 | 3 |  |
| 4 | 7.33±0.33 | 4 |  |

The results are mean± standard error of 3 replicates. 90 explants were used for/during the experiment.
For each level of NAA concentration at pH 4 \*\* represents highly significant difference at $p < 0.001$ by Tukey's-HSD test.

Surprisingly, a higher concentration of NAA > 2mg/L led to callogenesis, but no significant rhizoids formation occurred at low pH. However, NAA at a concentration 0.5 or 2 mg/L with pH 4, incubation in the dark at 23°C±2 led to a maximum number of white rhizoids surrounded by many hairy root-like structures. Both cotyledonous and hypocotyl explant resulted in a significant induction of rhizoids, however, the former took less time for initiation. The average no. of rhizoids induced at pH 4 were 38.33 and 37.66 in a medium supplemented with 0.5 mg/L and 2 mg/L NAA respectively from cotyledons after 3 weeks of inoculation. Various levels of pH tested against the range of NAA (0.5 mg/L – 4.0 mg/L) clearly showed that only at certain threshold concertation, auxins at pH can effectively induce S.E (**Fig 3**). Other than pH 3 in which the medium failed to solidify and couldn’t support rhizoid induction (**Fig 2A**), the sequence of effective media pH to exhibit a substantial number of rhizoids and RTBs from both explants was 4.0 > 5.0 > 6.0 > 7.0 supplemented with NAA 0.5 or 2 mg/L

Rhizoids formed at pH4 were shifted to a medium supplemented with 5,10,15,20 mg/L TDZ and 5 mg BAP for RTBs induction. Rhizoids produced in dark conditions started the formation of rhizoid tubers (RTBs) when exposed to Light/Dark 16/8hrs photoperiod (80 μmolm^-2^s^-1^) cool white fluorescent lights after 12 days of inoculation on media supplemented with BAP and/or TDZ (**Table 4**). We found that RTBs were induced only on pH4 while no such structures were observed on pH5.8. Unlike rhizoids, tubers were seen to be induced in light only while rhizoids in dark incubation resulted in shooting rather than RTBs. The data generated indicate that pH4 is unique for the formation of both rhizoids and RTBs with a high number of tubers induced on media supplemented with two cytokinins analogues 5 mg/L BAP and/or 5 mg/L TDZ (**Table 4**). RTBs were further investigated for their regeneration capacity, consequently excised and placed on a shooting medium supplemented with 3/5 mg/L BAP and/or TDZ with pH4 and pH5.8. Thereafter, the induction of the shoot was exceptionally high on pH5.8 in comparison to shoot morphogenesis which was found to be trivial at a lower pH level (**Fig 5**). This suggests that a lower pH with auxins (NAA) is required for rhizoid induction in the dark.Cytokinins (TDZ/BAP) addition in media with lower pH under light conditions induced novel structures – rhizoid tubers (RTBs) in Tomato (**Fig 4** A2&B3). The low H+ concentration of growth media remained ubiquitous to tomato tissue culture in this study by formation of callus in less time. RTB embryoids induction represents a new approach for somatic embryogenesis (S.E) and thus a new regeneration system for fast and efficient propagation.

**Table 4.** Effect of TDZ and BAP on rhizoid tubers (RTBs) induction.

| Sr. | TDZ mg/L | BAP mg/L | Average number of RTBs per Ex-Plant | Induction time |
| --- | --- | --- | --- | --- |
| 1 | 5 | - | 47.75± 2.5** | 12 Days |
| 2 | 10 | - | 30.5± 3.4 | 12 Days |
| 3 | 15 | - | 18.25± 3.9 | 12 Days |
| 4 | 29 | - | 14± 4.1 | 12 Days |
| 5 | - | 5 | 44.5± 3.4** | 12 Days |

The results are mean ± standard error of 3 replicates. Total 90 rhizoids were used during the experiment. For each level of TDZ/BAP concentration at pH 4, \*\* represents highly significant difference at $p < 0.001$ by Tukey's-HSD test.

**Table 5.** Effect of different concentrations of BAP, NAA, IAA & KIN on percent shooting frequency and average shooting time in tomato.

| Treatment | BAP (mg/L) | NAA (mg/L) | IAA (mg/L) | KIN (mg/L) | % Shooting Mean<br>Riogrande | % Shooting Mean<br>Roma | % Shooting Mean<br>Hybrid | % Shooting Mean<br>M82 |
| --- | --- | --- | --- | --- | --- | --- | --- | --- |
| SIMT1 | 2 | --- | --- | --- | 37.333±1.21<sup>e</sup> | 33.000±1.52<sup>e</sup> | 25.000±1.52<sup>c</sup> | 9.267±0.63<sup>b</sup> |
| SIMT2 | 3 | --- | --- | --- | 47.667±1.45±<sup>c</sup> | 40.000±1.15<sup>d</sup> | 40.000±2.84<sup>a</sup> | 7.667±0.48<sup>d</sup> |
| SIMT3 | 5 | --- | --- | --- | 54.667±2.40<sup>b</sup> | 63.333±1.14±<sup>a</sup> | 41.667±1.85<sup>a</sup> | 13.500±2.56<sup>a</sup> |
| SIMT4 | 1 |  | 0.1 | 2 | 44.667±1.76<sup>d</sup> | 28.000±1.76<sup>de</sup> | 25.667±2.33<sup>c</sup> | 5.200±0.6<sup>e</sup> |
| SIMT5 | 3 | 0.1 | --- | --- | 55.667±1.45<sup>b</sup> | 54.667±1.15<sup>c</sup> | 23.000±2.08<sup>c</sup> | 8.100±0.2<sup>c</sup> |
| SIMT6 | 3 | --- | 0.1 | --- | 69.000±2.08<sup>a</sup> | 58.667±1.45<sup>b</sup> | 32.33 ±2.96<sup>b</sup> | 5.600±0.4<sup>e</sup> |

Values are means of three replicates, the values bearing different letters in the same column are significantly different from each other according to the Fisher's least significant difference (LSD) test at p<0.001
SIM: Shoot induction medium

**Fig 4.**
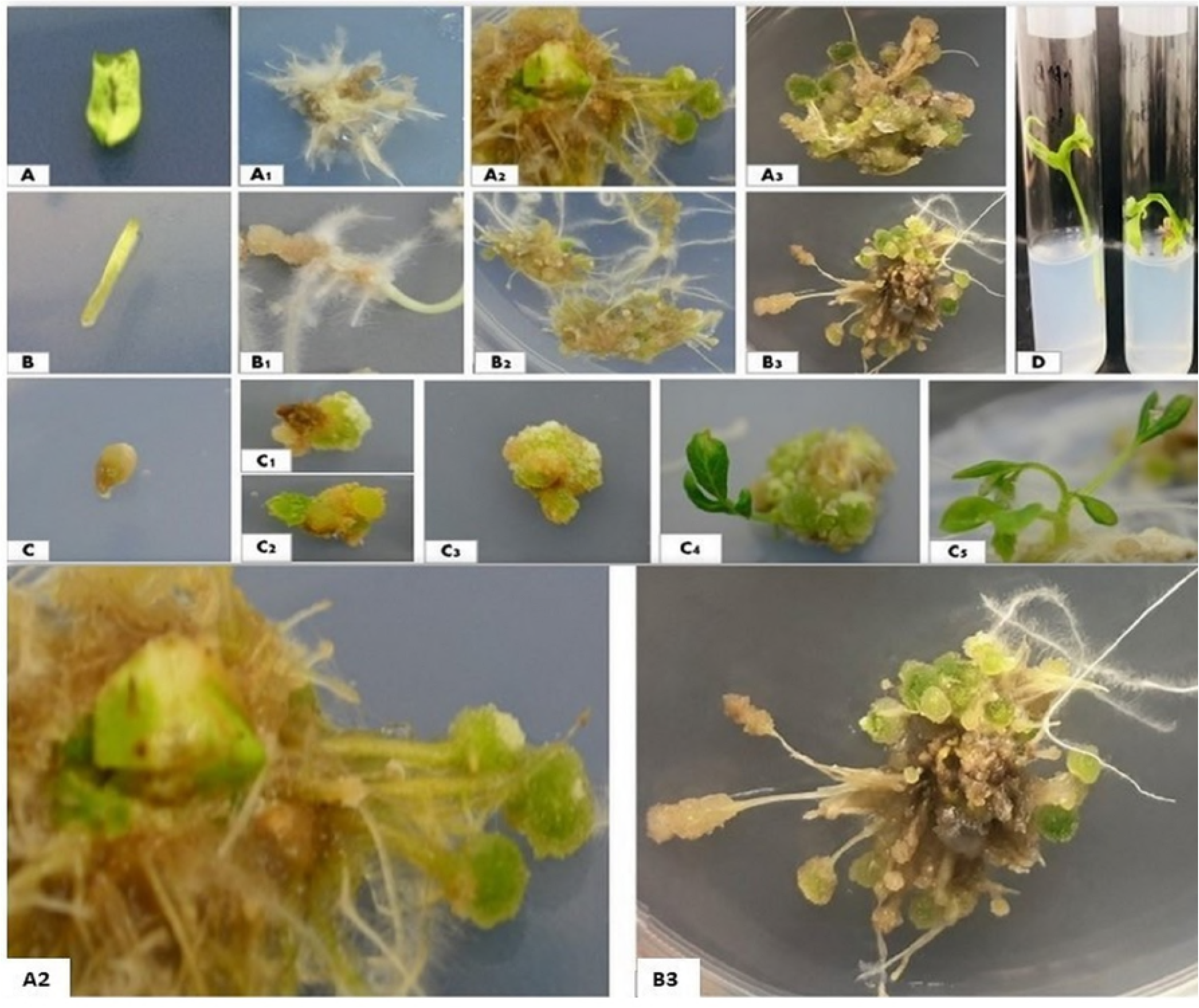
Development of somatic embryogenesis from leaf and hypocotyl of Tomato. (A-A3) Induction of rhizoids and rhizoid tubers from leaf, (B-B3) Hypocotyls, (C-C5) Steps of invitro somatic embryogenesis from individual Rhizoid tuber.

**Fig 5.**
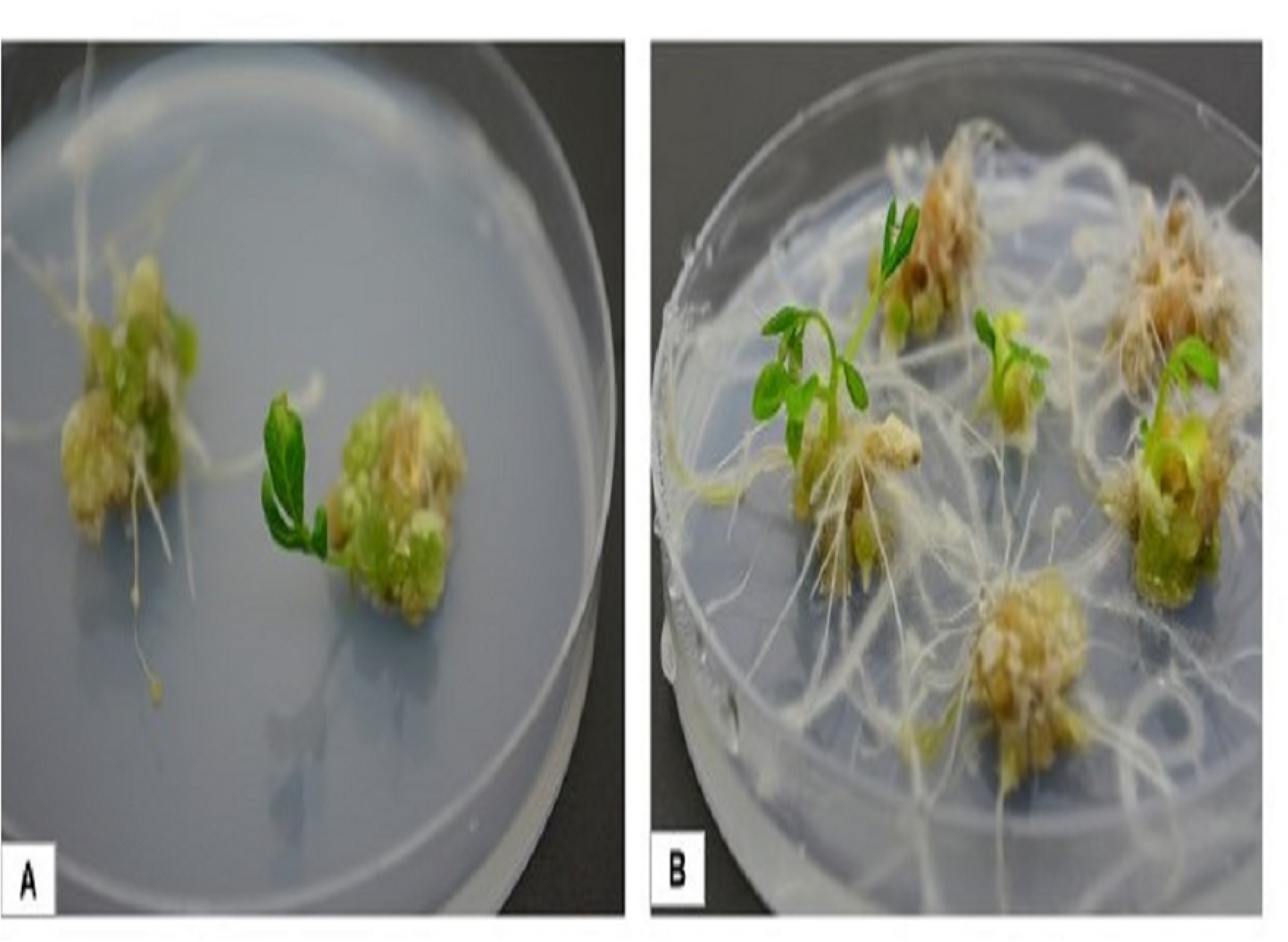
(A) Shoot and (B) Root emergence from individual rhizoid tubers. Significant differences were observed among treatments at (p<0.001); all four cultivars depicted differential response to each treatment. The order of shoot-induction frequency from cotyledons was Rio>Roma>Hybrid-17905>M82 (**Table 5**). The effect of treatment*genotype was found highly significant (**Table 5**). The time for morphogenesis of shoots ranged from 3.5 to 6 weeks after the inoculation on shoot induction media. The newly regenerated shoots were rooted on root induction media (RIM) containing different hormones [**Table 7**]. Rooting was observed 2 weeks post inoculation. NAA at 0.1 mg/L and 0.5 mg/L rendered a maximum number of roots. Excised shoots were also allowed to develop roots on MS media with and without PGRs. The addition of auxins NAA (0.1 mg/L) or rooting hormone IBA (1 mg/L) led to an abundant root formation (**Fig 6F**).

**Fig 6.**
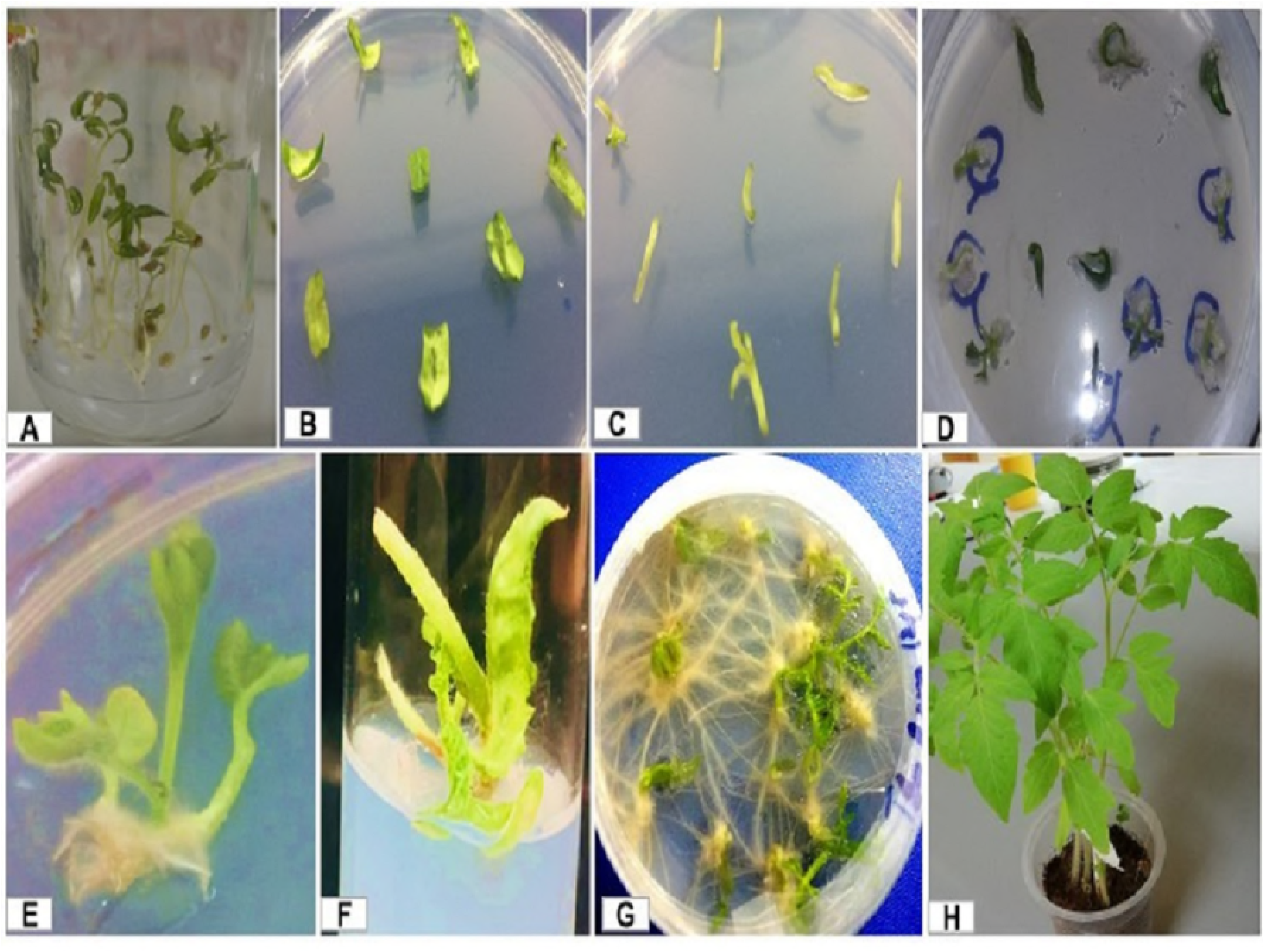
Stages of invitro morphogenesis from cotyledonous- and hypocotyl-based explants of *Riogrande*. (A) Germination, (B, C) Pre-culture treatment, (D) Callus induction, (E, F) Shoot induction, (G) Root induction, (H) Acclimatization.

### Whole plantlet regeneration from RTBs

Cotyledon and hypocotyl explants which developed compact green calli on all tested media were sub-cultured to the shoot induction medium after 4 weeks (**Table 5**). Rhizoid tubers were also inoculated on the shoot induction medium at pH4 and 5.8. Maximum number of shoots were induced on SIMT_6_ media with BAP (3 mg/L) and IAA (0.1 mg/L) with 6–7 shoots produced per explant. BAP alone at 3–5 mg/L was the second most effective medium (SIMT_3_ and SIMT_6_) in shoot regeneration. The shoot induction frequency was maximum for *cv. Riogrande* with 68.9% using SIMT_6_ from cotyledon-derived explants followed by RTBs. The result suggested that RTBs-derived shooting occurs in less time and is often accompanied with rooting (**Fig 5**). The regeneration results mentioned above had given rigorous shoot formation at pH5.8.

**Table 6.**
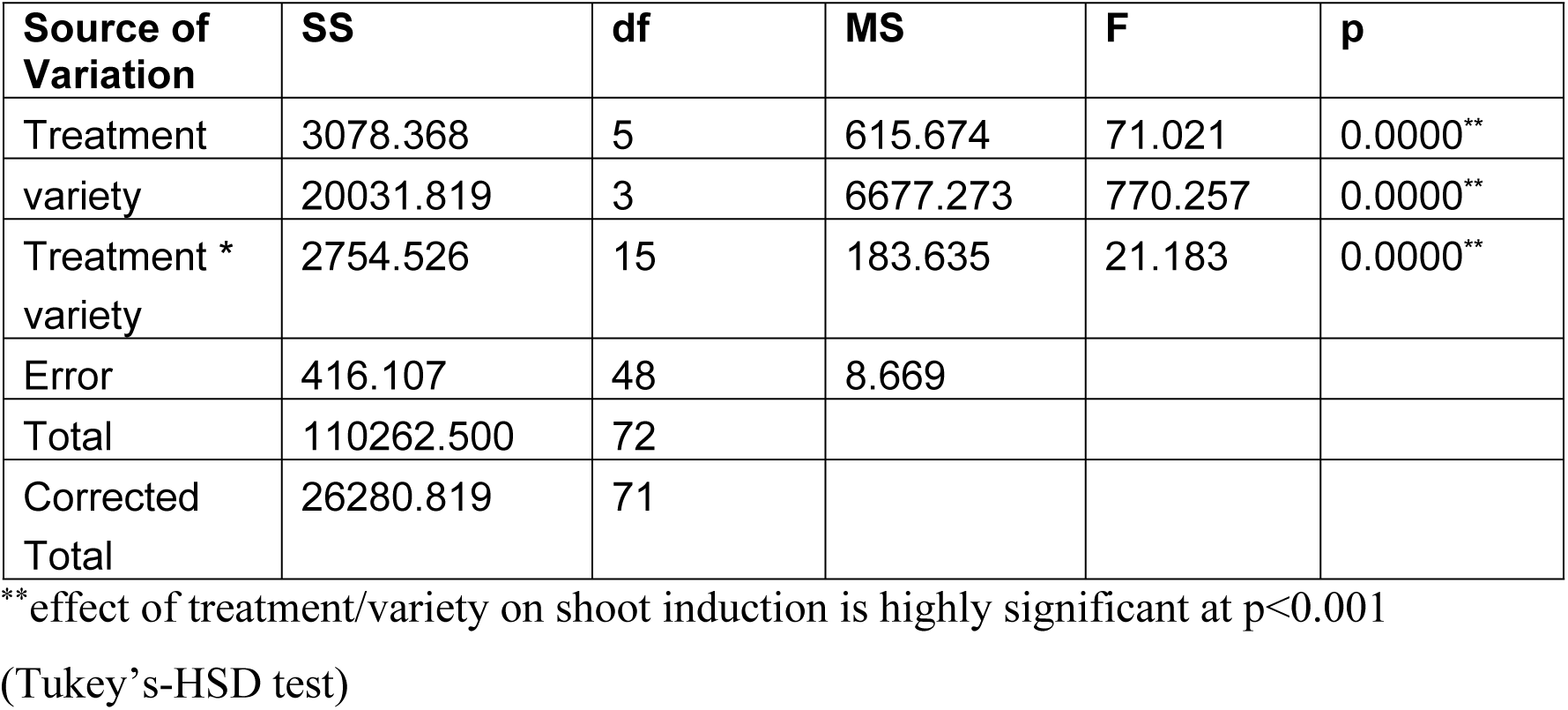
Analysis of variance for effect of treatment*variety on shoot induction in tomato

**Table 7.** Effect of different concentrations of IBA, NAA & IAA on average No. of roots produced from individual shoots in tomato after 8-10 weeks of incubation.

| Treatment | IBA (mg/L) | NAA (mg/L) | IAA (mg/L) | Mean No. of Roots<br>Riogrande | Mean No. of Roots<br>Roma | Mean No. of Roots<br>Hybrid | Mean No. of Roots<br>M82 |
| --- | --- | --- | --- | --- | --- | --- | --- |
| RIMT1 | 0.1 | --- | --- | 6.16±0.76<sup>d</sup> | 4.66±0.57<sup>c</sup> | 3.33±0.57<sup>c</sup> | 1.33±0.77<sup>b</sup> |
| RIMT2 | 0.2 | --- | --- | 4.66±2.08<sup>e</sup> | 5.33±1.15<sup>b</sup> | 2.33±0.57<sup>d</sup> | 0.33±0.57<sup>d</sup> |
| RIMT3 | 0.5 | --- | --- | 8±0.50<sup>c</sup> | 4.66±1.52<sup>c</sup> | 4±1.00<sup>b</sup> | 1.66±1.5<sup>b</sup> |
| RIMT4 | 1 | --- | --- | 9.66±1.52<sup>b</sup> | 6.33±1.52<sup>d</sup> | 2±1.00<sup>e</sup> | 4±1.00<sup>a</sup> |
| RIMT5 | --- | 0.5/0.1 | --- | 12.66±2.08<sup>a</sup> | 10.33±1.52<sup>a</sup> | 7.33±0.57<sup>a</sup> | 1±1.00<sup>c</sup> |
| RIMT6 | --- | --- | 0.1 | 7±1.00<sup>c</sup> | 5.66±2.08<sup>b</sup> | 4.66±1.52<sup>b</sup> | 1.33±1.52<sup>b</sup> |
| RIMT7 |  |  | 0.2 | 6.66±1.52<sup>d</sup> | 4.66±2.08<sup>c</sup> | 2.33±0.57<sup>d</sup> | 1.33±0.57<sup>b</sup> |

Values are means of three replicates, the values bearing different letters in the same column are significantly different from each other according to the Fisher's least significant difference (LSD) test at $p<0.001$
RIM: Root induction medium

### Histological Analysis of RTBs revealed proembryos and multiple embryoids

Microscopic observation on the transverse section of mature RTBs stained with safranin showed an internal arrangement of embryonic cells and non-embryonic cells. After 12 days of cultivation on the tubers induction medium (TIM), rhizoid pro-embryos turned visibly stained, dark pink while non-embryonic tissues were unstained (**Fig 7 A-D**). Further incubation led to the formation of RTBs embryoids with globular, heart, and heart-torpedo shapes (**Fig 8 A-E**). Each tuber exhibited multiple embryoids at different stages of development, hence progressing like typical somatic embryogenesis to a whole plantlet (**Fig 7 A&B**).

**Fig 7.**
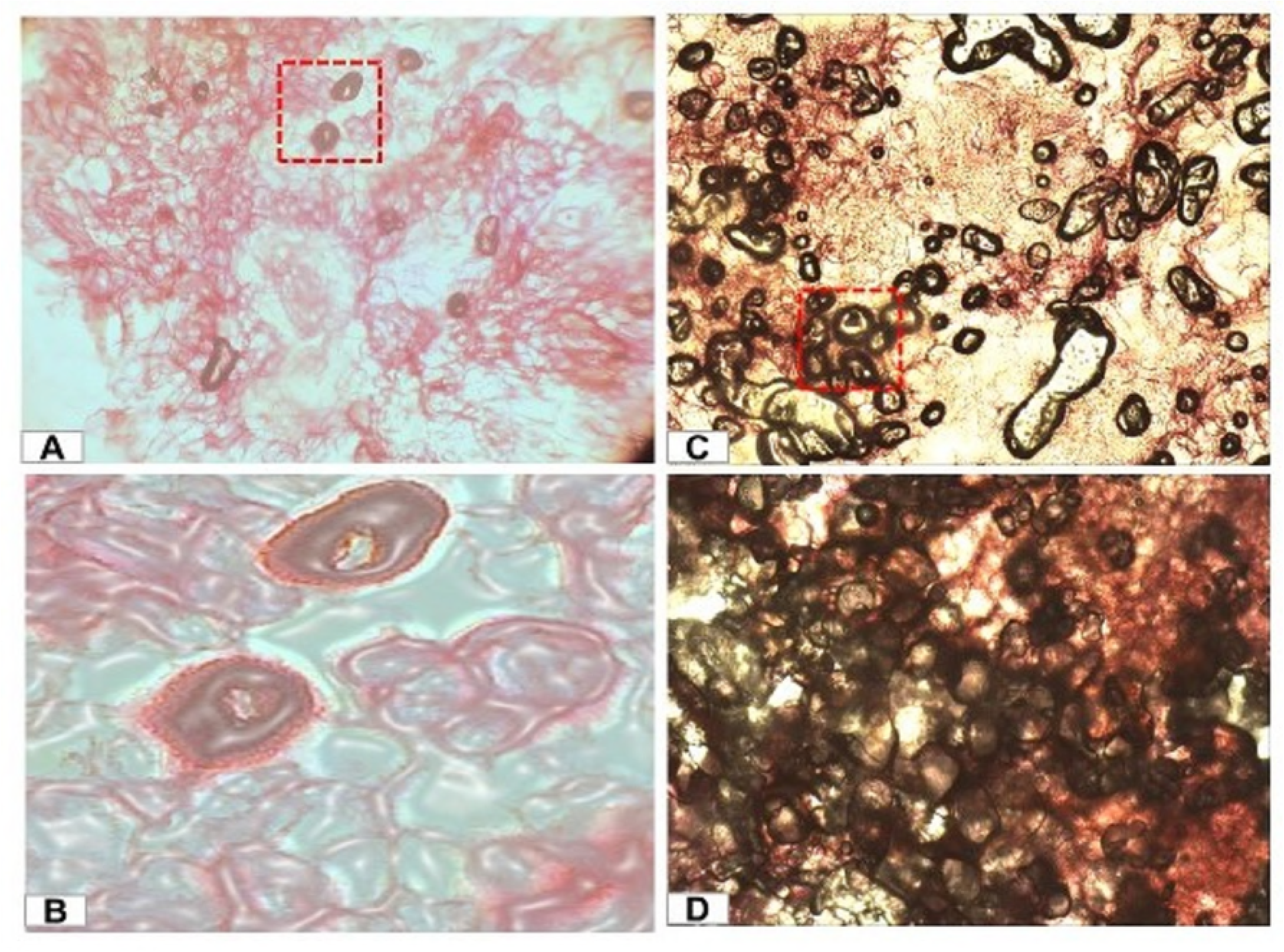
Histological study of mature Rhizoid Tubers (RTBs) of cv. *Riogrande* stained with safranin stain. (A) Transverse section of regenerating rhizoid tubers after 25 days of incubation under compound microscope [40X] 125μm, (B) Enlarged view of T-section showing embryonic cells, (C) Transverse section of mature RTB under 20X (100μm) automatic scanning system [ASS], (D) Enlarged view of section under box showing multiple embryoids.

**Fig 8.**
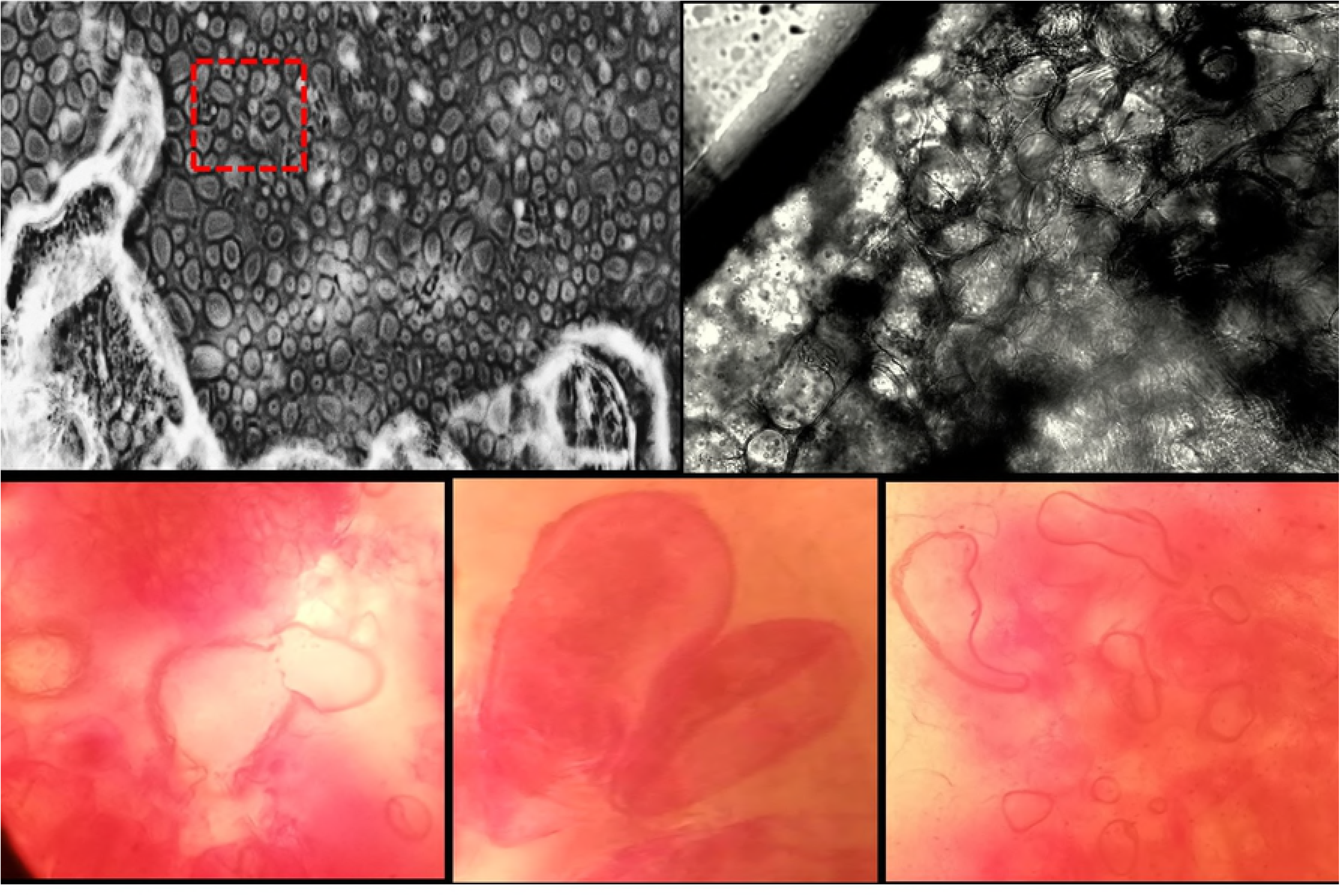
Direct somatic embryogenesis from cotyledons in cv *Riogrande*. (A) Morphology and arrangement of embryogenic cells under 20X (100μm) Automatic staining system (ASS) stained with Safranin, (B) Enlarged view of transverse section under box, (C) Globular shaped cells [nucleus not stained], (D) Heart shaped cells, (E) Torpedo-shaped cells.

## Discussion

Conventional breeding management augmented with sustainable *invitro* cell culture can be exploited for the production of genetically engineered plants. The development of the aforesaid system will be of great value in germplasm conservation, somatic embryogenesis, and the multiplication of *invitro* grown seedlings.

This research focuses on concurrent comparison of *in vitro* morphogenesis in tomato via organogenesis through callus induction and direct somatic embryogenesis (S.E). For this purpose, we optimized the germination and sterilization protocols for four different cultivars of tomato to increase their efficiency for *invitro* regeneration. *Invitro* morphogenesis response of the cultivars relies on genotype X environmental factors, thus making tomato transformation complex. Different concentrations of PGRs were used alone and in combination for a short-time induction organogenesis in all four cultivars. Explants (6 days post germination) were used for callus induction on 12 different media compositions which showed that higher cytokinins with low auxins tend to induce callus formation. Highest callus induction was observed from the treatment CIMT_7_ in both young cotyledons and hypocotyls of *Riogrande* and *Roma* where callus induction index was >80%. While *M82* and hybrid line 17905 responded to a completely different set of PGRs owing to genotypic differences. Such observations are in line with previous reports, where various combinations of PGRs were found cultivar-dependent and accounted as a trivial cause of slow *invitro* regeneration. Our results are consistent with [15,24] who reported that the regeneration capacity of explants decreased with an increase in age.

Embryogenic calli derived from leafy explants depicted a high-shoot regeneration potential in Rio and Roma while the other two cultivars had pale white compact calluses/calli (**Fig 1**). Hypocotyl derived callus has many embryoids (**Fig 1 A&B**) in *Riogrande*, however, callus induction was slower and calli were less totipotent as compared to cotyledons. Regenerating calli led to multiple shoot formations in a 3-week time period on an optimal shoot induction medium (SIM) containing more cytokinins (BAP 3 mg/L). On an average, 5-6 shoots were induced on SIM in case of Rio *cv*. with 69% shooting frequency. Previous studies have shown that high cytokinins favor short-time shoot multiplication in tomato in 13 different tomato cultivars [25,26]. In our study, the whole organogenesis process completed in less than 2.5 months for *Riogrande* with 3 weeks of callogenesis, 3 weeks for shoot multiplication, and 12 days of rooting. This high frequency standardized procedure yields acclimatized plantlets in 2-3 months.

We have developed organogenesis via S.E in *cv. Riogrande*. of tomato to further shorten the time of regeneration. The effect of medium pH on S.E is reported here as a novel aspect of tomato regeneration which influences the overall progress of clonal propagation and cell cultures. Another novel feature of the study is the development of S.E in tomato through rhizoid and rhizoid tubers. The results showed a quick regeneration response on pH4 in the presence of auxins leading to. S.E in hypocotyls and cotyledons of *Riogrande* (**Figs 2 and 3**) in two steps supplemented with NAA and TDZ respectively. In the first step, rhizoids are induced form young explants by using two auxin analogues at various concentration (NAA & 2,4-D) in dark conditions (**Fig 4 A1&B1**). NAA concentration at 0. 5-2 mg/L led to a successful rhizoid induction while 2,4-D did not induce these structures. This is the first report describing different auxinmediated responses for S.E in tomato. Individual rhizoids, upon transfer to a medium containing TDZ/BAP in light, produced novel structures – rhizoid tubers (RTBs) with many somatic embryoids (**Fig 4, A2&A3, B2&B3**). Different novel structures like RTBs, frog- egg-, and bulbil-like bodies have been reported through S.E recently [27–30] and our findings are in agreement with the previous reports [28]. Light conditions also played a significant role in the S.E of tomato. Dark conditions favored an induction of rhizoids while light enhanced RTBs formation. However, studies have also reported the positive effect of light treatment on S.E of different plant species [31,32]. During S.E, MS medium supplemented with a high level of auxins (2 mg/L NAA) and low pH, not only limited yeast contamination but also shortened the time of somatic embryogenesis and regeneration. A significant number of rhizoids was produced on MS medium containing NAA at pH4 while higher pH values resulted in callogenesis rather than rhizoids induction (Fig 4). To our knowledge, there is no report of *invitro* regeneration in tomato at low pH. However, low pH has been used to grow cell cultures of other plants. An increased No. of regenerated shoots at lower pH (4.5) in *Bacopa* monnieri has been reported [33]. pH range (3-7) was used to evaluate the regeneration capacity in pine buds showing that initially low pH of media is more suitable for the invitro morphogenesis [34]. The RTBs were further tested for their regeneration ability on SIM and RIM. Surprisingly, RTBs showed significant shoot and root formation ability at pH 5.8 in comparison to a lower pH (Fig 5 A&B). These results further demonstrate the superiority of novel structures - RTBs for complete organogenesis in less time and efforts over routinely used explants/sources. Histological studies of RTBs demonstrated their composition of embryogenic cells with multiple embryonic stages (**Figs 7 and 8**). These embryos sprout from rhizoids and then develop into whole plantlets through various growth stages.

Our findings suggest that the selection of right auxins at low pH (i.e. 4) along with alternate incubation in light/dark conditions are favorable conditions for the induction of S.E. The pH values < 4 were unable to support S.E while > 4 led to callus induction. These rhizoids were first induced from the cotyledon explants in the dark, which later developed into rhizoid tubers in light conditions via S.E. The presence of embryonic cells inside individual RTBs indicate their ability to form multiple embryos. RTBs can spontaneously regenerate into whole plantlets contrary to typical S.E. The organogenesis via S.E and tuber-shaped structure represents a robust way of a whole plantlet development in *Riogrande* and can be utilized for the establishment of similar systems in other recalcitrant cultivars of tomato like M82. Additionally, dedicated studies can be conducted in the future to use RTBs for the genetic transformation of desired traits in elite transgenic plants.

## Acknowledgement

The authors are thankful to the Institute of Agri-Biotechnology and Genetic Resources (IABGR), NARC, Islamabad for the provision of tomato seeds. The English language and grammar improvement of the manuscript has been done by Dr. Ayesha Siddiqa, Assistant Professor from Linguistic Department Quaid-i-Azam University Islamabad which is acknowledged.

## Authors Contributions

WS conducted the practical work, compiled the literature, and wrote the manuscript. DG helped WS in the practical work. SN helped in the execution of the research work, data compilation, and the revision of the final manuscript. ZA designed the strategy, supervised the overall research work, and improved the manuscript.

## Competing interests

The authors declare that the research was conducted in the absence of any commercial or financial relationships that could be construed as a potential conflict of interest.

## Funding

WS is highly thankful to the Higher Education Commission (HEC), Pakistan for providing financial support under the HEC Indigenous Ph.D. Scholarship Program (2BM2-161).

